# Early afterdepolarisation tendency as a simulated pro-arrhythmic risk indicator

**DOI:** 10.1101/176560

**Authors:** Beth McMillan, David J. Gavaghan, Gary R. Mirams

## Abstract

Drug-induced Torsades de Pointes (TdP) arrhythmia is of major interest in predictive toxicology. Drugs which cause TdP block the hERG cardiac potassium channel. However, not all drugs that block hERG cause TdP. As such, further understanding of the mechanistic route to TdP is needed. Early afterdepolarisations (EADs) are a cell-level phenomenon in which the membrane of a cardiac cell depolarises a second time before repolarisation, and EADs are seen in hearts during TdP. Therefore, we propose a method of predicting TdP using induced EADs combined with multiple ion channel block in simulations using biophysically-based mathematical models of human ventricular cell electrophysiology. EADs were induced in cardiac action potential models using interventions based on diseases that are known to cause EADs, including: increasing the conduction of the L-type calcium channel, decreasing the conduction of the hERG channel, and shifting the inactivation curve of the fast sodium channel. The threshold of intervention that was required to cause an EAD was used to classify drugs into clinical risk categories. The metric that used L-type calcium induced EADs was the most accurate of the EAD metrics at classifying drugs into the correct risk categories, and increased in accuracy when combined with action potential duration measurements. The EAD metrics were all more accurate than hERG block alone, but not as predictive as simpler measures such as simulated action potential duration. This may be because different routes to EADs represent risk well for different patient subgroups, something that is difficult to assess at present.

## Introduction

Torsades de Pointes (TdP) is a particular type of polymorphic ventricular tachycardia, characterised by an unusual electrocardiogram, in which the QRS complex appears to be twisted around the baseline. TdP usually spontaneously resolves, sometimes causing syncope (sudden fainting due to a drop in blood pressure), but it can also cause cardiac arrest or sudden death. [1, 2]

Anti-arrhythmic drugs, such as quinidine, [3] are commonly linked to TdP. Terfenadine, a non-sedating anti-histamine, was one of the first non-cardiac drugs to be linked to increased TdP risk. [4] Terfenadine was withdrawn from the market in 1997 after being linked to 41 cases of TdP, one of which was lethal. [5] Cisapride is a drug that was used for treating gastroesophageal reflux disease. [6] After causing 97 cases of TdP, of which six were fatal, cisapride was withdrawn from the market. [7]

Quinidine, terfenadine, and cisapride were all found to strongly block I_Kr_, the rapid delayed rectifying potassium current in the heart, which is carried by the channel whose primary subunit is a product of the human ether-a-go-go related gene (hERG). [8, 9, 4, 6]

Since these discoveries, testing for hERG block has become a mandatory requirement for new phar-maceuticals. [10] hERG block as a measure of TdP risk is very sensitive (gives few false negatives) and has prevented torsadogenic drugs from entering the market. Certain marketed drugs, such as ver-apamil and ranolazine, block hERG but are not linked with TdP. [11, 12] There are therefore concerns that hERG block lacks specificity (gives false positives for TdP risk), preventing the development of potentially useful drugs. [13] As such, elucidation of other factors that mediate TdP risk is needed.

Early afterdepolarisations (EADs), phenomena in which the membrane depolarises a second time during the action potential, are heavily implicated in the onset of TdP. [15] Hearts suffering from TdP show EADs alongside transmural dispersion of repolarisation. [16, 17, 18, 19] EADs are seen in monophasic action potential recordings from dog hearts during TdP, as shown in Figure 1. [19, 14]

**Figure 1:**
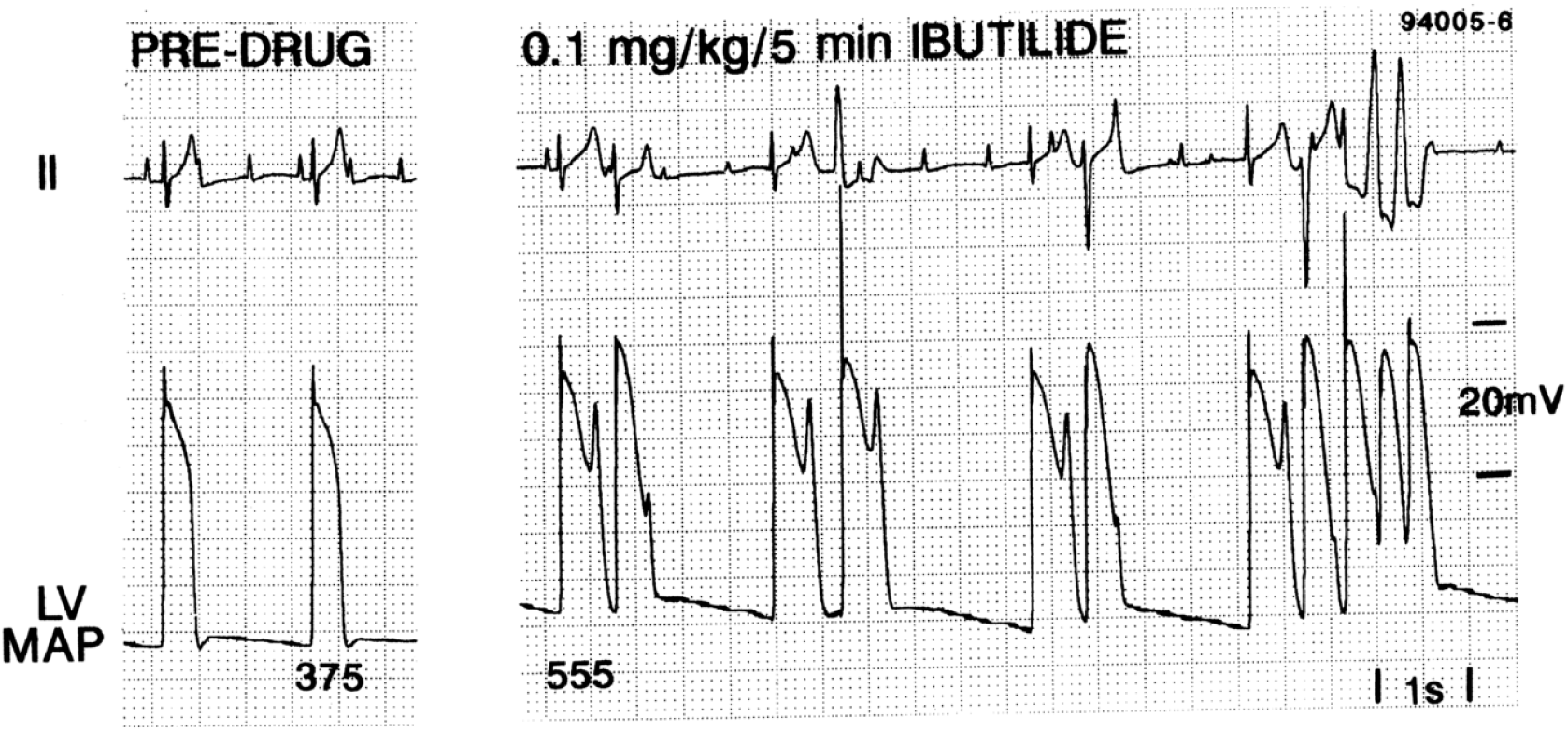
Induction of EADs and TdP in an anaesthetised dog by treatment with ibutilide [14]. Reproduced with permission from Paul G. A. Volders *et al*. “Progress in the understanding of cardiac early after depolarizations and torsades de pointes: time to revise current concepts”. Cardiovascular Research (2000) **46**(3): 376–392. Published by Oxford University Press on behalf of the European Society of Cardiology (ESC).

Our investigation is based on several mechanisms that are known to promote EADs. One mechanism is based on a form of Long QT syndrome (LQT8) that is caused by an increase in the L-type calcium current, caused by a gain of function mutation in the CACNA1C gene, that increases the density of L-type calcium channels at the cell surface up to three-fold. [20, 21] Patients with LQT8 frequently exhibit TdP. [22] L-type calcium current agonists (agents which increase the conductance of the L-type calcium channel) have also been shown to cause: TdP in live mice[23]; EADs and ventricular tachycardia in intact mouse hearts[24]; EADs in sheep and dog Purkinje fibres[25]; and EADs in ferret ventricular myocytes[26]. L-type calcium current blockers such as verapamil are used as anti-arrhythmic agents, and have been shown to suppress EADs and TdP in rabbit models of LQT3 and chronic heart failure. [27, 11, 28]

Another EAD mechanism we investigate in this study is based on Brugada syndrome, which is a condition linked to ventricular fibrillation and elevation in the ST segment of the electrocardiogram. [29] Estimates of the proportion of sudden deaths caused by Brugada vary from 4–12%, and it is estimated to be present in 0.05% of the world population. In 15–20% of Brugada cases, the cause is a mutation in the SCN5A gene, which codes for the alpha subunit of the fast sodium channel. [29] A missense mutant version of SCN5A (T1620M), was shown to alter the inactivation curve of the fast sodium current, shifting it in the positive direction by 10 mV [30], and another missense mutant, L812Q, shifted the inactivation curve by 20 mV. [31] Adding this inactivation curve shift to cardiac action potential models can cause EADs (Noble *et al.*, personal communication).

In addition to being of interest in drug-induced TdP, the I_Kr_ potassium current is also involved in LQT2, and is also linked to EADs [9]. Loss-of-function mutations in hERG cause a reduction in the I_Kr_ current of up to 97%,[32] which causes QT prolongation, and usually increases risk of TdP. LQT mutations do not always present with significant clinical QT prolongation, but interaction with drugs that affect cardiac ion channels could cause an increased risk of TdP. [33]

Two other Long QT conditions that are linked to TdP are LQT1 and LQT3. [34] LQT1 causes an increase in the slow delayed potassium current I_Ks_, and LQT3 causes an increase in the persistent sodium current I_pNa_. We have not used these as EAD-provoking interventions in this work because we were unable to provoke EADs in the O’Hara cell model by blocking I_Ks_ or increasing I_pNa_.

The prediction of arrhythmias by *in silico* modelling of action potentials in response to ion channel block offers a new way to test novel compounds at the pre-clinical stage. A previous study by our group created an improved measure of a compound’s propensity for causing TdP arrhythmias, using simulated action potential duration as a metric. [35] The approach takes into account the contributions of multiple ion channels to the shape and length of the action potential, and classifies drugs into discrete risk categories, based on their effect on action potential duration. This method was more accurate than the commonly-used ‘hERG safety factor’, that is the ratio of hERG IC_50_ to effective free therapeutic plasma concentration (EFTPC), or log_10_(hERG IC_50_*/*EFTPC_max_). [36] Simulation studies have been extended to predict results of rabbit wedge studies and the Thorough QT study. [37, 38]

A recent study used principal component analysis to assemble a large number of biomarkers from different models, the results suggested that a two-dimensional binary classification based on both the simulated diastolic calcium concentration and the APD50 was effective at separating drugs into positive or negative for torsadogenicity. [39] We aim to extend these approaches to allow prediction of the TdP risk classes of drugs that increase action potential duration but are safe (particularly late/persistent sodium blockers), and to account for the interaction of drug block with underlying conditions such as ion channel mutations. The appearance of EADs at increased drug concentrations has been studied computationally as a risk indicator for TdP. [40] Our study complements this work by exploring EADs as a risk indicator in the context of disease states at clinically-relevant concentrations.

Ion channel conductance modification as a cause of EADs has also been investigated recently in the context of atrial fibrillation: a global sensitivity analysis of atrial cell models was used to examine which ion channel changes lead to EADs and then logistic regression was used to estimate the probability of EADs as functions of conductances. [41]

The Comprehensive in-vitro Pro-arrhythmia Assay (CiPA) is a proposal to use multi-ion channel screening in combination with human stem cell-derived cardiomyocytes and computational cardiac modelling to create new metrics for the prediction of drug-induced TdP, moving away from using QT interval prolongation as a surrogate marker and towards *in vitro* and *in silico* methods. [42] This study investigates whether a marker linked mechanistically to EAD formation may be helpful in determining torsadogenic risk.

We applied ion channel block to cardiac cell models to simulate the effects of drugs of known torsadogenic risk, then combined these drug effects with simulated disease states. The level of disease state necessary to provoke an EAD for each drug was used as a marker for TdP risk. These markers were evaluated against clinical risk categories, separately, combined, and in combination with pre-existing TdP risk metrics. By combining these markers we hope to create a measure of risk for a population that contains people with each EAD-inducing condition. In this way, we bring together three key factors that influence arrhythmogenic risk: the ion channel blocking properties of the compounds, the concentrations of compound found in humans, and underlying risk factors from the patient.

## Methods

We selected 41 drugs of known torsadogenic risk, and simulated their ion channel blocking effects in the O’Hara 2011 human ventricular cell model.[43] Using a range of interventions, we determined the threshold of intervention at which an EAD could be provoked in the cell model, i.e. the lowest level of intervention that was necessary for an EAD to be produced. The differences in ‘threshold for EAD’ between different drugs were used to classify drugs by arrhythmic risk. These steps are detailed below.

### Drug inclusion criteria

To select drugs to use as a training set, we used three criteria based on the amount of data available on both the pro-arrhythmic risk of the compound and its effect on ion currents in cardiac cells.

1. As a starting point, we included drugs that we previously studied in Mirams et al. [35].
2. Additional drugs were then included in our study if they had been included in five or more of the papers analysing TdP risk discussed in a recent summary paper, [44] and if over 70% of studies agreed on high or low TdP risk.
3. Drugs that are on the CiPA list [42] were automatically included if ion current block data were available for three or more channels of interest (even if they had been in fewer than five studies or had poor agreement in risk category between studies).

If there was disagreement in risk class using the above sources, the default category was the one used in Mirams et al. [35], and if not listed there then the one used in Redfern et al. [36].

To be included in our dataset, the drugs were also required to have IC_50_ values available in the literature for three or more of the ionic currents of interest, found by manual patch clamp. The ionic currents of interest were: the fast and late/persistent sodium currents, the L-type calcium current, the rapid and slow delayed rectifier potassium currents, the transient outward current, and the inward rectifier potassium current. The pIC_50_ values we used are given in Table 1, and the references for these can be found in the Supplementary Material. Where there were multiple IC_50_ values available, preference was given to those from human cells at body temperature (37°C) and to the most recent study.

**Table 1:**
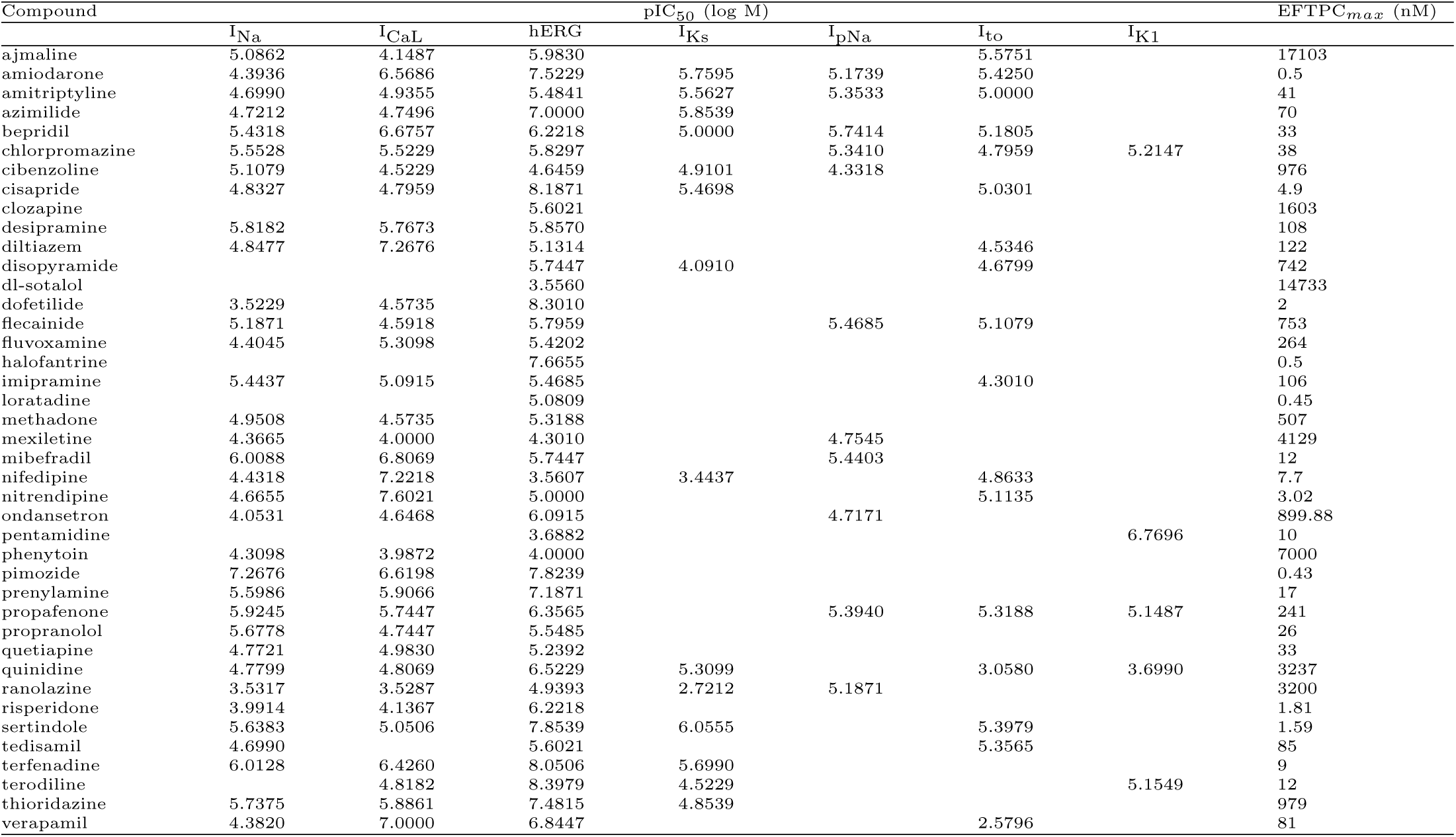
pIC50 values for each compound in the dataset (shown as log M) for the fast sodium (I_Na_), L-type calcium (I_CaL_), rapid delayed rectifier potassium (hERG or I_Kr_), slow delayed rectifier potassium (I_Ks_), persistent sodium (I_pNa_), transient outward (Ito) and inward rectifier potassium (I_K1_) currents, and effective free therapeutic plasma concentration (EFTPC) (nM). For references, please see the Supplementary Material. ‘n/a’ indicates that the channel has been screened and no effect was measured: either the IC50 value was above the maximum concentration being tested, or there is no drug-induced block of this channel.

Stratification into 4 risk categories was taken from the Mirams *et al*. (2011) paper where available, and the Redfern *et al.* (2003) paper otherwise. We pooled risk categories 1 and 2 together, as they represent the same risk level for different drug classes, leaving classes 2–5. [35, 36] The only compounds in the dataset not covered by this metric were: (i) ranolazine, which was assigned to Category 4, i.e. the drug has isolated reports of TdP in humans; and (ii) cibenzoline, which was assigned to Category 5 based on Lawrence et al. [45], i.e. there are no reports of TdP in humans with this drug. [46, 47]

### Action potential and drug block models

We used two recent human ventricular myocyte electrophysiology models based on human datasets, the O’Hara (2011)[43] endocardial and Grandi (2010)[49] models. Drug block of ion channels was modelled as a reduction in channel conductance as a function of the concentration of the compound, [*D*], and the IC_50_ value.[50] The change in maximum conductance for a channel *j* was described by a Hill equation:

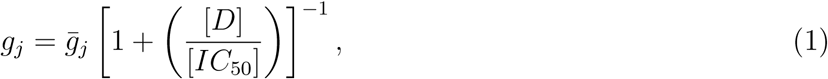

where *g*_*j*_ is the maximum conductance of the drug-blocked channel, and 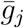 is the conductance of the channel when there is no compound present. The Hill coefficient here is set to 1, as the variability in experimentally-inferred Hill coefficients from patch clamp can be so high that using 1 as the Hill coefficient may reduce error [51]. Note that this formula was applied for all channel/drug combinations listed in Table 1, apart from late/persistent sodium in the Grandi model — as this model does not have a distinct late/persistent sodium current.

Drug concentrations were set to the maximum effective free therapeutic plasma concentrations (EFTPC) for each individual drug, to provide a realistic estimate of ion channel block *in vivo*. For a list of EFTPCs, see Table 1.

### Detecting afterdepolarisations

The appearance of early afterdepolarisations (EADs) in a simulation was determined by the slope of the voltage trace between adjacent time points. Whenever the slope was greater than +1 mVms^−1^, a depolarisation was reported. To remove depolarisations caused by the stimulus, depolarisations that occurred between 50 ms before and 100 ms after each stimulus were disregarded. The algorithm detects single EADs, multiple EADs, and EADs without repolarisation, as shown in Figure 2. For the full algorithm, see the `DetectAfterDepolarisations` class in the code repository (see “Numerical methods and simulation procedure”).

**Figure 2:**
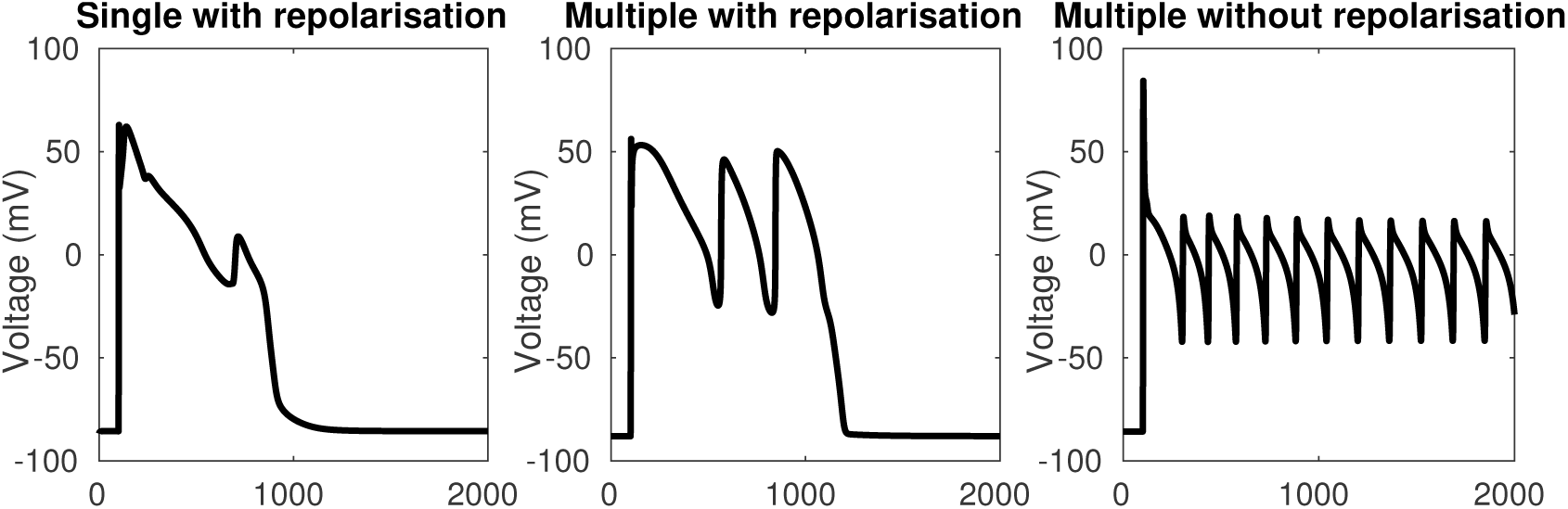
Types of EADs that are detected using the EAD detection algorithm. Left: a single EAD with repolarisation. Centre: multiple EADs with repolarisation. Right: multiple EADs without repolarisation.

### Provoking afterdepolarisations

The failure of hERG block alone to predict torsadogenic risk suggests that drug-induced TdP is mediated by more than one ionic mechanism. We hypothesise that the interaction of certain disease states with torsadogenic drugs could lead to greater susceptibility to TdP. We propose to look at the single-cell phenomenon of EADs, as studies suggest TdP and EADs are intrinsically linked.[16, 17, 18, 19, 14] Spatial differences in ion channel expression throughout cardiac tissue would alter the susceptibility of each cell to EADs, meaning that some regions would have EADs while others did not, increasing the probability of re-entrant waves, but we constrain this study to looking at the ‘trigger’ event of EAD initiation.

To simulate disease states after drug block was applied, EADs were provoked using three interventions mimicking disease states discussed in the introduction, based on a manuscript from Noble et al. [52]:

- conductance of the L-type calcium current was increased by a scaling factor to simulate a gain-of-function mutation which causes LQT8. [21]
- the midpoint of the inactivation curve of the fast sodium current was shifted using an additive constant to simulate Brugada syndrome. [30]
- conductance of the rapid delayed rectifier current I_Kr_ was decreased by a scaling factor to simulate the effect of LQT2 mutations. [9]

The O’Hara et al. [43] model was used for these EAD simulations, as it includes all of the currents of interest, and is being investigated for CiPA-related proarrythmic risk prediction. [42]

We used a slow pacing interval of 3 seconds because bradycardia is also known to facilitate onset of EADs and TdP. [17] We performed the same simulations at faster pacing rates, which did not significantly change the classification results.

### Previously suggested measures

In order to compare our EAD tendency measures with previously suggested risk indicators, we used our dataset of drug actions to calculate some risk metrics that have previously been proposed.

We calculated the hERG IC_50_*/*EFTPC_max_ safety margin proposed by Redfern et al. [36]; the predicted APD_90_ calculated using the Grandi et al. [49] human ventricular cell model, as in Mirams et al. [35]; the predicted APD50 combined with diastolic calcium concentration calculated using both the Grandi 2010 Grandi et al. [49] model, and the O’Hara 2011 [43] model as in Lancaster and Sobie [39]; and the predicted sum of normalised total persistent sodium current and L-type calcium current over the course of the action potential at increasing drug concentrations (known as ‘cqInward’) using the O’Hara et al. [43] model, as in Li et al. [48], but using the baseline model rather than their dynamic hERG block model (as we did not have hERG kinetic data for all compounds). The Grandi model was chosen for APD_90_ rather than the O’Hara model as it was used in the Mirams et al. (2011) metric.

### Numerical methods and simulation procedure

Simulations were run using the Chaste C++ framework [53, 54] with the ApPredict bolt-on project [55] and custom written code. Cell models and initial conditions were imported from CellML [56] files using PyCML [57].

The adaptive timestep solver CVODE [58] was used to solve the model’s differential equations, with a relative tolerance of 10^−5^ and absolute tolerance of 10^−7^. The output timestep was 0.1 ms and a stimulus current was applied every 3 s for 3 ms with a magnitude of −25.5 *μ*A*μ*F^−1^.

The procedure we used is outlined in Algorithm 1. First, drug block was applied to the model by modifying the conductance parameters for the appropriate channels, using Equation (1) to calculate updated conductances. The model was then run to steady state using the pacing protocol above. Steady state was reached when the norm of the change in the model’s state variables was less than 10^−6^ between paces. This generally took 1,000–10,000 paces.

EAD-provoking interventions were applied by modifying either the I_CaL_ conductance, the I_Kr_ conductance, or the I_Na_ shift parameter. After setting the intervention parameter, the simulation was run for 12 s, and the presence or absence of EADs was detected. Interval bisection was used to find the threshold of intervention that was necessary to cause any EADs. We started with interval ranges of: [0, 20] mV for the I_Na_ shift intervention; [1, 80] for the I_CaL_ conductance scaling; and [0, 1] for the I_Kr_ conductance scaling. Interval bisection was set to terminate when consecutive values were less than 10^−4^ units apart. EAD simulations for a single compound can be run in less than 10 minutes, and many compounds can be run in parallel. All the code is available to download from github at https://github.com/teraspawn/EadPredict.

#### Algorithm 1 procedure to find the EAD threshold

1: Set ion channel conductances in cell model to new values based on drug block.

2: Run cell model to steady state and then save this state to reset to later.

3: Set upper and lower limits *α, β* for intervention value.

4: **repeat**

5: Reset cell model to earlier state.

6: Set intervention to (*α* + *β*) *÷* 2 (e.g. multiply I_CaL_ conductance by this factor).

7: Run model for 12 s.

8: Check for Early afterdepolarisation (upwards trajectory after initial depolarisation).

9: Adjust *α* or *β* using interval bisection.

10: **until** *|α - β| <* 10^−4^

### Linkage analysis

To visualize whether drugs with similar risk require similar EAD thresholds, linkage analysis was used to create dendrograms of drug similarity based on each of the metrics. The Euclidean distance between the metric values (e.g. APD_90_ for each of the drugs) was used to construct a dendrogram, grouping drugs which had similar values. To combine metrics, the results were shifted such that the control value was zero, and then scaled to be within [*-*1, 1]. Combinations of metrics such as all the EAD metrics, the EAD metrics with APD_90_, and all the metrics together, were then used to create classification trees. The optimal leaf ordering was calculated using the R package “dendextend”, to best sort the leaves in descending order of risk category (without changing the branching structure).[59]

This method allows for the grouping of drugs by similarity rather than by rigid categories, allowing for new compounds to be visually ranked by closeness to torsadogenic and non-torsadogenic drugs. The output of the optimal leaf ordering algorithm was evaluated by the sum of the square difference between the risk category of each drug in the ranking and an optimal ordering (2,2,…,2,3,3,…,3,4,4,… etc.):

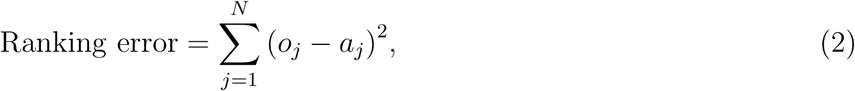

where *o* is the optimal ordering of risk category, *a* is the actual risk category, and *N* is the number of drugs in the dataset.

### Pro-arrhythmic risk classification

As in Mirams et al. [35], risk categories were based on the Redfern et al. [36] classes (with Category 1 and Category 2 pooled, as they represent equivalent levels of risk for different drug classes):

- Category 2: Either Class Ia and III antiarrhythmics, or drugs that have been withdrawn from market due to TdP.
- Category 3: drugs with a measurable incidence or numerous reports of TdP in humans.
- Category 4: drugs with isolated reports of TdP in humans.
- Category 5: drugs with no reports of TdP in humans.

We also considered a binary classifiers, where Categories 2 and 3 were grouped as torsadogenic and Categories 4 and 5 were grouped as non-torsadogenic. Results from this were not materially different, and can be found in the supplementary spreadsheet under “Binary”.

We used EAD thresholds, APDs, diastolic calcium concentration, and hERG IC_50_*/*EFTPC_max_ to classify compounds into one of the four TdP risk categories described earlier. We tested two classification methods: linear discriminant analysis (LDA), and support vector machines (SVM).

Leave-one-out cross-validation and five-group cross-validation were used to check the robustness and accuracy of both the classifiers. We evaluated performance by calculating the errors in classification (how many risk classes away from the correct class a drug was classified as), and comparing these means of absolute error, *E*:

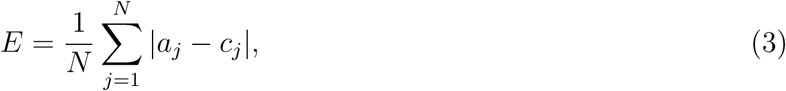

where, *a* is the actual category (i.e. the real risk category for the drug), *c* is the category assigned using the classification method, and *N* is the number of drugs in the dataset.

In LDA, the metrics in categories are assumed to follow a Normal distribution, and the points where the inferred distributions overlap are used as the category boundaries.[60] In SVM, a hyperplane is used to separate the data points, and the optimal hyperplane is found by maximising the distance between the hyperplane and the closest points to it.[61]

In leave-one-out cross-validation one drug was removed from the dataset and the boundaries were re-calculated. The left-out drug was then placed into a risk category based on these new boundaries. 5-group cross-validation was also used: drugs were randomly assigned to five groups, and then each group was removed from the dataset and the classification boundaries were re-calculated. The drugs in the removed group were then classified into a risk category based on these new boundaries, and accuracy was calculated as above.

## Results

### EAD thresholds

In general, more torsadogenic drugs caused a decrease in the threshold of intervention required to provoke an EAD, i.e. they made the electrophysiology models more vulnerable to EADs. Full tables of EAD thresholds can be found in the Supplementary Material.

Some examples of EADs induced by the I_CaL_ increase protocol are shown in Figure 3. The low risk drug, nitrendipine, required a greater increase in I_CaL_ conductance (27.23*×*) than the control (24.13*×*) to induce an EAD. Conversely, the high risk drug cisapride required a much smaller increase (8.23*×*) to induce an EAD. Similarly, models whose ion channel conductances had been modified to simulate the effects of cisapride required an I_Na_ inactivation curve shift of only 15.45 mV to produce an EAD, whereas at control a larger shift of 17.41 mV was required, and the low risk drug diltiazem required a shift of 17.49 mV, greater than control.

**Figure 3:**
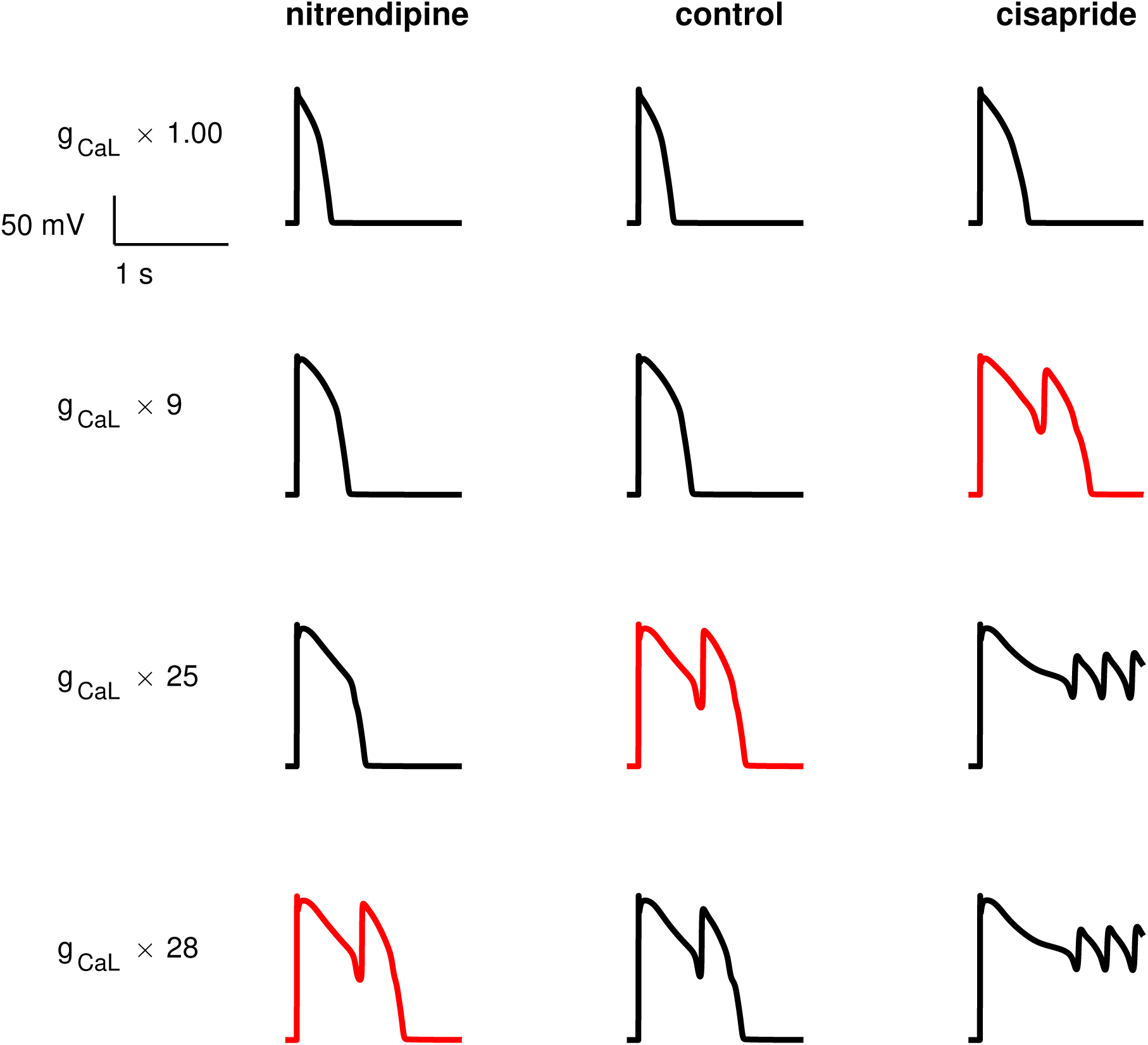
Some examples of EADs provoked using the L-type calcium increase protocol, as described in the methods section. The first EAD caused by the increasing intervention for each drug is highlighted in red. The less torsadogenic drug (nitrendipine) requires more provocation than the control to cause an EAD (i.e. its EAD threshold is higher), and the more torsadogenic drug (cisapride) requires less provocation (i.e. its EAD threshold is lower).

In general, I_CaL_-provoked EAD thresholds were lower for drugs which strongly block the I_Na_ and hERG channels. I_CaL_ EAD thresholds were not linear with I_CaL_ drug block, showing that multi-ion channel effects affected the thresholds. Less I_Kr_ block was required to cause an EAD for strong hERG blockers, except for verapamil, which is a very strong hERG blocker, but also strongly blocks I_CaL_.

Thioridazine needed no hERG block to cause an EAD, despite being a strong I_CaL_ blocker. This is probably due to thioridazine’s very strong effect on hERG, with an IC_50_ value of 0.034*×*EFTPC_max_.

I_Na_ inactivation curve shift thresholds were lower for drugs which strongly block hERG. Unlike the other EAD metrics, block of I_CaL_ and I_Na_ did not increase the EAD threshold for the I_Na_ inactivation curve shift protocol. For example, verapamil, which is a strong hERG and I_CaL_ blocker, had a low I_Na_ shift EAD threshold.

All three of the EAD metrics consistently had the lowest thresholds for the following drugs: azimilide, cisapride, ondansetron, terfenadine, and terodiline. Three drugs (ajmaline, quinidine, and thioridazine) produced EADs without any intervention required. With the exception of thioridazine (category 3), all of these drugs are in the highest risk category 2.

A table of EAD thresholds and other metrics can be found in the Supplementary Material.

### Linkage analysis

The dendrogram from the linkage analysis of the hERG IC_50_*/*EFTPC_max_ metric, APD_90_, and the combined EAD metrics are shown in Figure 4. The optimal leaf ordering algorithm was moderately successful in sorting the drugs in ascending order of torsadogenic risk. Therefore, a classification scheme based on the similarity of new compounds to existing compounds of known torsadogenic risk could be a useful tool. The ranking error measure for all metrics can be found in the Supplementary Material.

**Figure 4:**
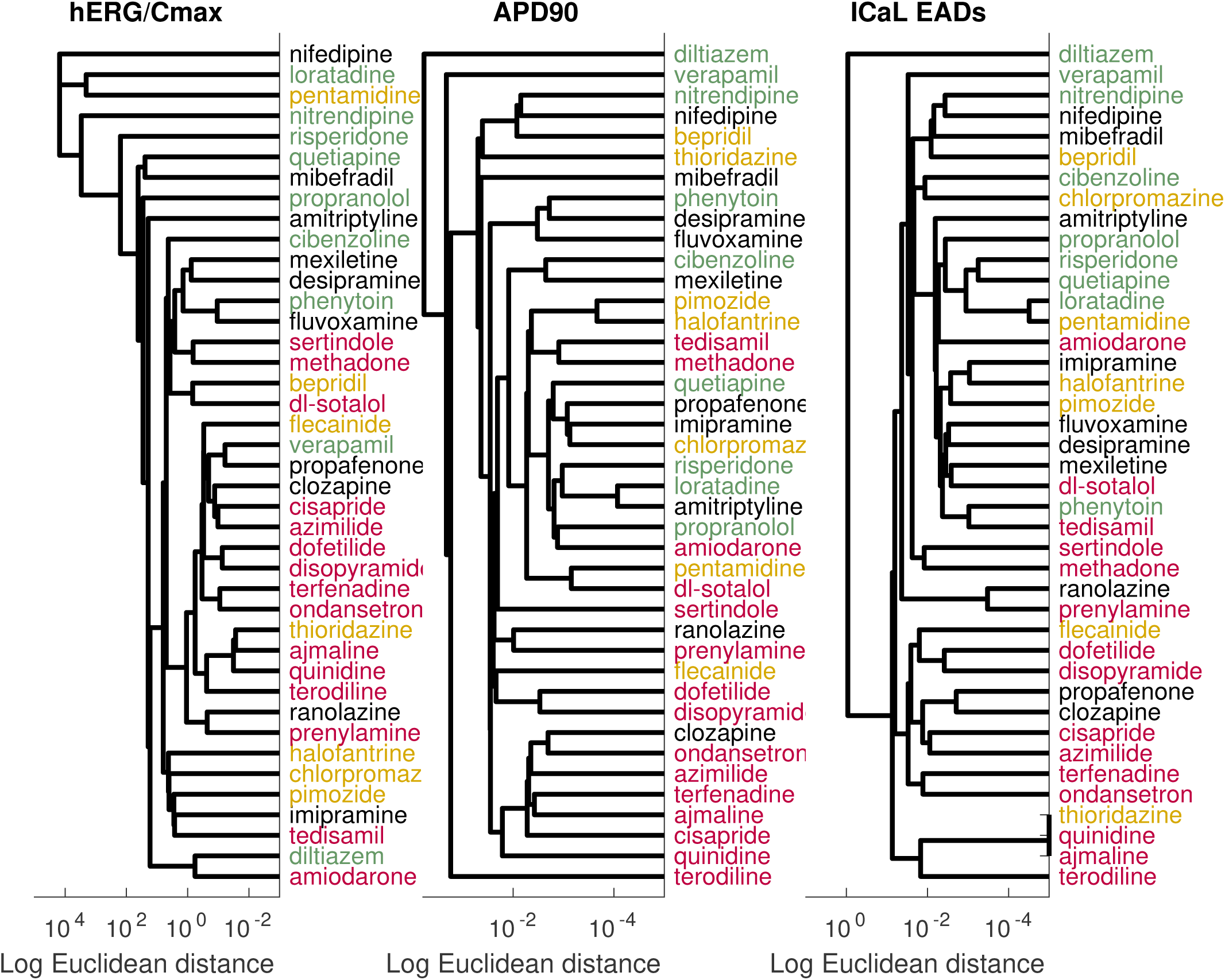
Left: linkage analysis based on the hERG IC_50_*/*EFTPC_max_ metric. Centre: linkage analysis based on the APD90 metric. Right: linkage analysis based on the I_CaL_ EAD threshold metric. Risk categories are indicated by colour: Category 2 (dangerous) drugs are shown in red, Category 3 in yellow, Category 4 in black, and Category 5 (safe) in green.

### Pro-arrhythmic risk classification

Table 2 shows the mean absolute error, *E*, in classification for each of the proarrythymic risk markers and classification methods. Overall, there was good agreement between the different classification schemes and validation procedures. Full tables of all results can be found in the Supplementary Material. The leave-one-out analysis showed that the Support Vector Machine method of classification was more accurate for the Grandi Lancaster-Sobie metric, the Grandi APD_90_ metric, the O’Hara Cqinward metric, and the hERG IC_50_*/*EFTPC_max_ metric. For all other metrics, Linear Discriminant Analysis was more accurate.

**Table 2:** Mean absolute errors in classification, calculated using Equation 3. The “5 LDA” and “5 SVM” rows are the results from the 5-group cross-validation of linear discriminant analysis classification and support vector machines classification, respectively. The “1 LDA” and “1 SVM” rows are the results from leave-one-out cross-validation. The columns are arranged by the lowest sum of errors from the four classification measures. Yellow cells show metrics with low errors and purple cells have high errors. ‘L&S’ refers to the diastolic calcium concentration-APD50 combination metric from a paper by Lancaster & Sobie [39], ‘cqInward’ refers to the metric from Li et al. [48] and ‘Redfern hERG Safety Factor’ refers to the hERG IC_50_*/*EFTPC_max_ metric proposed in Redfern et al. [36], as discussed above.

| Classifier | Grandi<br>APD<sub>90</sub><br>+<br>O’Hara<br>I<sub>CaL</sub> | Grandi<br>APD<sub>90</sub> | Grandi<br>L&S | O’Hara<br>I<sub>CaL</sub><br>EADs | Grandi<br>APD<sub>90</sub><br>& all<br>O’Hara<br>EADs | All<br>O’Hara<br>EADs | Redfern<br>hERG<br>Safety<br>Fac-<br>tor | O’Hara<br>L&S | O’Hara<br>I<sub>Na</sub><br>EADs | O’Hara<br>cqIn-<br>ward | O’Hara<br>I<sub>Kr</sub><br>EADs |
| --- | --- | --- | --- | --- | --- | --- | --- | --- | --- | --- | --- |
| LDA 1 | 0.78 | 0.80 | 0.95 | 0.78 | 0.95 | 1.00 | 1.17 | 0.95 | 1.05 | 1.20 | 0.90 |
| SVM 1 | 0.78 | 0.71 | 0.68 | 1.07 | 0.98 | 1.17 | 0.85 | 1.07 | 1.20 | 1.10 | 1.29 |
| LDA 5 | 0.83 | 0.98 | 0.93 | 0.93 | 0.88 | 1.00 | 1.17 | 1.27 | 1.07 | 1.02 | 1.00 |
| SVM 5 | 0.90 | 0.90 | 0.88 | 0.85 | 0.85 | 1.00 | 0.98 | 1.02 | 1.00 | 1.05 | 1.66 |

Using LDA, the most accurate metrics were the O’Hara I_CaL_ increase EAD metric and the O’Hara I_CaL_ increase EAD metric combined with the Grandi APD_90_ metric. Using SVM, the most accurate metric was the Grandi Lancaster-Sobie metric, followed by Grandi APD_90_.

The least accurate metric using LDA was the cqInward metric. Using SVM, the least accurate metric was the I_Kr_ EAD protocol, followed by the I_Na_ inactivation curve shift EAD metric.

The best metric over all classification methods was the Grandi Lancaster-Sobie metric using SVM, and the worst over all classification method was the I_Kr_ EAD protocol using SVM.

The I_CaL_ increase EAD metric correctly classified cibenzoline, desipramine, and fluvoxamine into Categories 5, 4, and 4, respectively, while the APD_90_ metric mis-classified them into the more dangerous Categories 4, 3, and 3, respectively, and the hERG IC_50_ / EFTPC_max_ metric mis-classified all three drugs into the most dangerous Category 2.

## Discussion

In this paper, we have presented a novel method for predicting pro-arrhythmic risk by provoking EADs in computational models of cardiac cells, in combination with simulated ion channel block.

As expected, EAD thresholds were usually lower for drugs which strongly block hERG, except when hERG block was combined with other ion channel block. I_CaL_ block removed the effect of hERG block on the I_Kr_ EAD threshold, for example, verapamil, a strong I_CaL_ blocker, required a large decrease in I_Kr_ current to cause an EAD despite being a strong hERG blocker. This indicates that for patients with LQT2 or other hERG mutations, drugs which block I_CaL_ could be beneficial in preventing TdP. I_CaL_ EAD thresholds were lower for drugs which strongly block both the I_Na_ and hERG channels. Therefore, for patients with increased I_CaL_ activity, due to I_CaL_ agonists or genetic mutations, drugs which block both I_Na_ and hERG could increase the risk of TdP. I_Na_ inactivation curve shift thresholds were not increased by I_CaL_ block — for example, verapamil had a low I_Na_ EAD threshold despite strongly blocking I_CaL_. This indicates that, for patients with Brugada syndrome, the torsadogenic effects of hERG block cannot be ameliorated by I_CaL_ block. I_CaL_ increase as a torsadogenic risk metric showed clear improvement over the hERG ‘safety margin’ marker of IC_50_ / EFTPC_max_.

In general, results were consistent over classification methods and evaluation scheme, as shown by leave-one-out cross-validation and five-group cross-validation.

The I_CaL_ increase EAD metric sorted cibenzoline, desipramine, and fluvoxamine into their correct risk categories, while both hERG IC_50_ / EFTPC_max_ and Grandi APD_90_ classified them into more dangerous categories. All three of these drugs are relatively strong I_CaL_ blockers. Cibenzoline is also a strong I_Na_ blocker and a weak I_pNa_ blocker. Block of the persistent sodium or L-type calcium currents have been shown to suppress EADs. [62, 63] These results show that EAD metrics can accurately predict the torsadogenicity of drugs which block hERG and increase APD without causing TdP.

Amiodarone was persistently misclassified by every metric except for hERG IC_50_*/*EFTPC_max_ using LDA, and the I_Kr_ EAD metric and cqInward metric using SVM. Amiodarone weakly blocks several ion channels, but also has active metabolites that were not considered here. [64] This combination of ion channel block leads to only a small increase in APD, and slightly decreased EAD thresholds. Ion channel block by amiodarone has been shown to be use-dependent, and there is evidence that amiodarone binds only to the open state of the I_Na_ channel. Therefore, a more detailed model of amiodarone binding kinetics may be required. However, despite being a Class III antiarrythmic, amiodarone poses a much lower, if still measurable, TdP risk than most other compounds in risk category one [65] and may therefore be a candidate for re-classification into a lower clinical risk category.

Following APD-based metrics, an EAD-based approach to pro-arrhythmic risk prediction offers a mechanistic link between ion channel block and TdP. [35, 37] Building on previous EAD-based metrics, [40, 41] our method looks at causes other than increased drug concentration as a cause of TdP, and incorporates the action of underlying disease states. Linkage analysis may allow for more fine-grained risk ranking and be helpful in showing compounds with similar properties.

The separation of drugs into risk categories based on TdP clinical incidence is a difficult problem. TdP can only be diagnosed when a patient is being monitored on an electrocardiogram, so episodes of TdP may be missed, giving underestimates of incidence. In addition, there is little publicly-available information on drug prescription numbers, meaning that it is difficult to calculate the number of TdP cases per prescription/dose. The lack of information about TdP incidence per dose makes risk classes uncertain, and a drug prescribed to people who are more likely to get electrocardiograms will have an increase in the number of cases of TdP diagnosed for that drug compared to a similarly torsadogenic drug. Wiśniowska and Polak [44] showed how different risk classifications have been given to the same drugs in different studies. Our strategy of using compounds with uncontroversial risk categorisations could ameliorate this problem, but the differences in risk category are fundamentally due to a lack of data on incidence.

The approach we have presented suggests a strategy by which risk assessments might be made for different patient subgroups (analogous to the different disease-mimicking EAD provoking interventions we applied). But as discussed, at present incidence data for the population as a whole is lacking, and this problem is exacerbated for smaller patient subgroups, making evaluation of our patient-group-specific predictions impossible at present. To improve these and other TdP prediction efforts, more data on TdP incidence rates will be needed.

One weakness of our study is the variety of sources from which the IC_50_ values were obtained. The differences in experimental protocol, temperatures, cell types and equipment used in these experiments may add uncertainty to our simulations, [66] which we have not included in the computations presented here. A dataset from a single set of manual patch-clamp experiments with low variability would allow standardisation across pro-arrhythmic risk classification methods for a range of drugs.

There are a large number of available cardiac cell models. For the afterdepolarization aspects of this study, we have used only the O’Hara et al. [43] model, because it includes the persistent sodium current. The addition of an appropriate late sodium current to existing models would be a useful extension to this work, to allow us to look at predictions from a wider range of models. This is work we are pursuing. We did not use the new dynamic hERG block model from Li et al. (2017) [48] which may explain why our implementation of the Cqinward metric was not as successful as in that paper.

Our study does not look at pro-arrhythmic markers at the tissue or whole heart level. EADs are a cell-level phenomenon that interact with several other factors in the onset of TdP in tissue. [67] The effects of spatial heterogeneity in ion channel expression and in fibre organisation on the arrhythmogenic effects of EAD susceptibility induced by drug block are likely to be significant in the translation from single-cell EADs to tissue- and organ-level effects.

Instead of using a simple pore-block model for drug interactions with ion channels, it would be interesting to account for different hERG binding kinetics, which have been explored in a recent study.[48] The differences in binding may alter the effect of the drugs on action potential duration and EAD susceptibility.

## Conclusions

We have proposed and investigated novel metrics for predicting drug-induced torsadogenic risk based on early-stage pre-clinical data on ion channel block. Our EAD-based metrics combine ion channel block data with disease states in order to predict increased EAD susceptibility in cell models, as a marker for pro-arrhythmic risk. Some EAD metrics were an improvement on the hERG IC_50_ / EFTPC_max_ safety margin as a predictor of clinical incidence of Torsades de Pointes. The I_CaL_ increase metric performed well, it more accurate than hERG block alone, but was not as predictive as simpler measures such as simulated action potential duration we have published previously. This may be because different routes to EADs mimic different diseases in patient subgroups and represent risk well for these patients only, but evaluating this is difficult without further data on clinical TdP incidence rates.

## Conflicts of Interest

The authors have no conflicts of interest to declare.

### Acknowledgements

Thanks to Martin Fink, Alan Garny, Penny Noble & Denis Noble for sharing their ideas on the generation of EADs and a paper draft. BM was funded by the EPSRC (Grant Number EP/G03706X/1). GRM gratefully acknowledges support from a Sir Henry Dale Fellowship jointly funded by the Wellcome Trust and the Royal Society (Grant Number 101222/Z/13/Z).

